# Decoding Cognitive States from fMRI Using Classical Machine Learning and Temporal Dynamics Analysis: An Interpretable Approach Using the Human Connectome Project

**DOI:** 10.64898/2026.05.29.728756

**Authors:** Classifying Brain States with ML Methods, Valeriya Kirova, Dzerassa Kadieva, Isak B. Blank, Daniil Vlasenko, Fedor Ratnikov

## Abstract

We propose a rigorous and reproducible methodology for analyzing functional MRI data, aimed at: (1) demonstrate their efficiency in classifying task-induced brain states with a limited amount of data, (2) present a methodology to identify brain regions critical for classification and reveal their uniqueness across different states, and (3) show, using strong mathematical methods, that the discriminative power of these regions depends not only on their spatial localization but also on their coordinated temporal activity. Through correlation and temporal structure analyses, we demonstrated that top-ranked regions exhibit stronger, more structured, and richer dependencies than low-ranked regions, underscoring the critical role of temporal dynamics in shaping distinct cognitive brain states. Our work addresses the need for a transparent, accessible, and interpretable framework for studying cognitive processes through neuroimaging data. We analyzed fMRI data from 587 healthy participants from the Human Connectome Project across seven cognitive tasks. Finally, we perform a detailed analysis of the identified brain regions to support further neuroscientific interpretation and discussion.

**Key Points:**

1. Classical machine learning methods effectively classify task-induced brain states from fMRI data with high accuracy (up to 99% for some tasks), demonstrating that simple, interpretable algorithms can successfully decode complex neuroimaging data without requiring advanced deep learning approaches.
2. High-accuracy brain states require relatively few significant regions suggesting focal neural signatures, while lower-accuracy states involve more distributed activations across multiple brain areas, revealing different levels of neural organization complexity underlying various cognitive processes.
3. The identified brain regions align with established neuroscientific knowledge, with motor tasks activating contralateral sensorimotor areas, language processing engaging left-hemisphere networks, and social cognition recruiting visual motion processing regions, validating the neurobiological relevance of our machine learning approach.
4. Rigorous mathematical analyses of temporal dynamics demonstrated that the discriminative power of significant brain regions depends not only on spatial localization but also on their coordinated temporal activity. Correlation, temporal structure analyses consistently showed that top-ranked regions exhibit stronger, more structured, and richer dependencies than low-ranked regions, underscoring the critical role of temporal dynamics in shaping distinct cognitive brain states.

## 1 Introduction

The brain contains specific regions and networks that play a key role in classifying and processing various states (emotional, cognitive, perceptual, etc.). It relies on a combination of activity across different regions and their interactions. Modern neuroimaging techniques, such as fMRI, enable the investigation of brain activity during these states, opening new avenues for studying cognitive processes. However, working with fMRI represents a complex challenge due to the high-dimensional and dynamic nature of the data. To address this challenge, we applied time-tested machine learning (ML) methods to fMRI data from the Human Connectome Project (HCP) (the 1200 Subject Release; Elam et al., 2021), aiming to efficiently classify different task-induced brain states.

In recent years, network-based data representation methods have gained increasing attention in neuroscience for describing functional connections between different brain regions (Wang et al., 2010; Richiardi et al., 2011; Takerkart et al., 2014). These methods model the brain as a complex network, where nodes represent brain regions and edges represent functional connections, enabling researchers to investigate both local interactions and global network topology (Li et al., 2021; Saeidi et al., 2022; Bessadok et al., 2023). Such approaches provide valuable insights into the organization and dynamics of functional brain networks, revealing patterns that underlie various cognitive states and neurological conditions. However, despite their growing popularity, network-based methods often require complex computations and substantial data preprocessing, which can limit their accessibility and scalability.

In turn, classical ML methods remain powerful tools for decoding fMRI data. These methods are generally less complex and computationally demanding, yet can achieve comparable classification performance by leveraging carefully selected regional activity features. Thus, while network-based analyses enrich our understanding of brain connectivity, they are not always necessary for effective classification.

Namely classical algorithms have delivered high results, including near-perfect identification of subjects and robust task-state decoding from the HCP data using off-the-shelf methods applied to functional connectivity features (Hannum et al., 2023). In their recent study with 865 participants and eight cognitive states, five standard classifiers — including support vector machines (SVM) — Hannum et al. (2023) reached accuracies up to 99%. This study shows that careful feature engineering and evaluation can be more efficient than complex and computationally expensive architectures.

Moreover, Popov et al. (2024) recently conducted a study comparing different ML classification models on fMRI data. The comparison revealed that model complexity does not guarantee better generalization in typical neuroimaging settings, especially with modest datasets, while the state-of-the-art accuracy can be achieved with relatively simple methods (Popov et al., 2024).

Beyond task decoding, resting-state analyses show that linear SVMs and related models can successfully separate meaningful stimuli-free brain conditions (Meier et al., 2019). For example, rs-fMRI amplitude metrics combined with sequential feature selection achieved roughly 81% accuracy distinguishing hunger from satiety, illustrating how straightforward pipelines can extract state-relevant signal without complex modeling.

Notably, interpretability remains a practical advantage of classical models: linear classifiers support direct inspection of weights to localize features and brain regions linked to each state, aiding neuroscientific inference alongside accuracy (Thapaliya et al., 2025). This transparency helps validate findings against prior knowledge and clarifies what aspects of the data drive predictions (Meier et al., 2019).

In our study, we build upon the technique proposed by Kirova V., Kadieva D., Ratnikov F. (2025), adapting it to our data and research goals. We first employed linear classifiers – such as logistic regression – which are widely used in neuroimaging research due to their robustness and interpretability. The classical methods have demonstrated successful applications in fMRI classification tasks, especially when data are limited and interpretability is crucial (Norman et al., 2006; Varoquaux, 2018). We refined this approach by identifying brain regions significant for classification – we ranked them according to their contribution, and iteratively removed the least influential regions. This recursive feature elimination and importance ranking based on classifier weights were used to reduce dimensionality and improve interpretability while maintaining classification accuracy in neuroimaging studies (Norman et al., 2006; Varoquaux, 2018). Furthermore, we tested whether dynamic temporal correlations contribute to functional connectivity beyond static averaged activity, by comparing correlation distributions of original and temporally shuffled time series. Numerous studies have shown that temporal variability and dynamic functional connectivity in fMRI reflect meaningful fluctuations in brain network organization relevant to cognition and pathology (Hutchison et al., 2013; Preti et al., 2017). Methods involving the comparison of original and temporally disrupted signals have been used to reveal the critical role of temporal structure in functional brain networks (Liégeois et al., 2019). Finally, we validated the results of this three-step process by comparing them with established neuroscientific findings to assess their accuracy and relevance.

Thus, the present approach achieves high accuracy with standard classifiers, but departs in three ways from existing studies applying ML algorithms on fMRI data classification. First, our approach prioritizes regional activation patterns over connectivity matrices. Second, it applies systematic feature selection to reduce to a minimal, informative set of regions. And third, it subjects models to explicit temporal shuffling to quantify the contribution of temporal structure.

Although each technique we selected for the current study is neither novel nor original on its own, together they provide a comprehensive and rigorous framework that can yield novel insights into the spatial specificity and temporal dynamics of task-induced brain states.

## 2 Methods and Materials

### 2.1 Dataset Description

The fMRI data we used for the ML classification was obtained from the Human Connectome Project (HCP) 1200 Subjects Release (Van Essen et al., 2013; Human Connectome Project, 2017) available through the MGH-USC HCP database (https://ida.loni.usc.edu/login.jsp).

Participants were selected using three inclusion criteria that ensured high level of quality of the HCP S1200 data. Only individuals who successfully completed all seven HCP task paradigms were retained. We have also excluded all cases listed as having quality issues during the HCP quality control process, retaining only participants with no reported imaging problems. Finally, we included only adults aged 22 to 35 years at the time of the scanning.

The final sample included 581 healthy participants, all right-handed (*n* of females = 295). Participants were met with a series of tasks (Barch et al., 2013) designed to assess seven main cognitive domains sampling the diversity of human neural networks, including: 1) visual, motion, somatosensory, and motor systems; 2) category specific representations; 3) working memory/cognitive control systems; 4) language processing (semantic and phonological processing); 5) social cognition (Theory of Mind); 6) relational processing; and 7) emotion processing. Three of these tasks were completed in one session, while the remaining four were performed in another session. A short description of these tasks is provided in Supplementary Material 1, for a more detailed description, see Barch et al. (2013). Each participant performed seven tasks with two conditions, resulting in 14 distinct brain states (Table 1) and 8134 fMRI data units in total.

**Table 1:** Two brain states for each cognitive task, between which classification is made.

| Cognitive Task | State name 1 | State name 2 |
| --- | --- | --- |
| Working Memory | 0-back | 2-back |
| Gambling | Win | Loss |
| Motor Task | Left hand or foot | Right hand or foot |
| Language Processing | Story | Math |
| Social Cognition | Random motion | Mental interaction |
| Relational Processing | Relation | Similarity |
| Emotion Processing | Neutral | Fear |

### 2.2 Data Acquisition

The data were acquired with a 32 channel head coil on a modified 3 T Siemens Skyra with TR = 720 ms, TE = 33.1 ms, flip angle = 52°, BW =2290 Hz/Px, in-plane FOV = 208 × 180 mm, 72 slices, 2.0 mm isotropic voxels, with a multi-band acceleration factor of 8 (Feinberg et al., 2010; Moeller et al., 2010). For our analysis, we only used the right-to-left phase encoding to maintain consistency. Data preprocessing followed the HCP minimal preprocessing pipeline, including motion correction, magnetic field distortion correction, spatial normalization, and filtering of spatial and temporal noise. Additional preprocessing steps were applied, including linear trend removal and z-score standardization of voxel-wise time series. For more details on data acquisition and preprocessing guidelines, see Barch et al. (2013) and Glasser et al. (2013).

Brain parcellation was performed using the HCP Multi-Modal Parcellation (MMP) atlas, which defines 379 distinct brain regions. This includes 180 cortical areas per hemisphere and 19 subcortical regions. For each region, representative time series were obtained by averaging the time series of all voxels within that region. This fine-grained parcellation enables high-resolution analysis of both functional connectivity and task-evoked activation patterns across the cortex and subcortex.

### 2.3 Data Structure

We defined the total number of brain states as *s* = 14 and the total number of brain regions as *r* = 379 . For each brain region, we have a time series of length *l* describing brain activity over the time interval.

Thus, each fMRI data unit is represented as an activity matrix **X** of size *r* × *l*, where each line 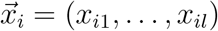 corresponds to the activity of the *i*-th brain region over a time interval of length *l*. Each element *x*_*ij*_ of the activity matrix **X** corresponds to the activity level of the *i*-th brain region at time point *j*.

We applied standardization to the data, transforming the values such that their distribution has a mean of 0 and a standard deviation of 1. We represented each activity matrix **X** by the averaged activity values across brain regions. For each brain region *i* = {1, …, *r*} we calculate the mean value 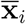 of the time series 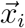. Thus, for each activity matrix **X** , we obtain a vector of mean activity values across all brain regions, which will be used in further analysis:

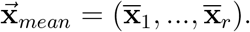

By focusing on the mean activity values, we aim to demonstrate that each brain state is associated with a unique network of regional activations, which can serve as a reliable signature for classification.

### 2.4 Classification Model

In ML formulation, each of the 379 averaged brain regions signals 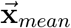 is considered as a *feature*, and the cognitive brain state to be predicted from fMRI data is considered as a *class label*.

Formally, we construct a multi-class classification model that maps a feature vector 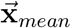, to a set of probabilities for that vector to represent each of *s* brain state classes:

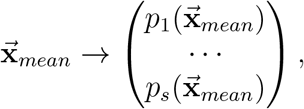

where 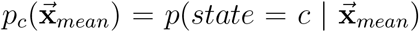, and *s* = 14 classes again correspond to distinct brain states. In simple words, each component 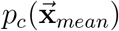 predicts the probability that input vector of mean activity values 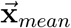 belongs to the brain state class *c*.

We begin with the simplest ML approach – linear models – to classify brain states, motivated by several key factors related to both our data characteristics and analytical goals. Their interpretability allows direct examination of weights highlighting important brain regions, while their efficiency with moderate-sized fMRI datasets helps prevent overfitting. These models demonstrate particular robustness to noise, maintaining reliable performance despite common artifacts in neuroimaging data. Their computational simplicity enables rapid processing of high-dimensional neural data without sacrificing predictive accuracy. Furthermore, linear models provide a flexible foundation that can be extended to capture more complex relationships when needed. This approach aligns well with established neuroscientific theory suggesting that cognitive processes emerge from linear combinations of regional brain activations.

We conducted a comprehensive analysis of linear ML models for multiclass classification of brain states using fMRI data. Our evaluation revealed that all linear models achieved high classification accuracy, with most models clustering around 0.9 accuracy as shown in Table 2. These results demonstrate that linear approaches retain competitive performance for brain state classification, suggesting their continued relevance in fMRI decoding tasks despite advances in nonlinear methods.

**Table 2:** Accuracy of linear classification models (ascending order)

| Model | Accuracy |
| --- | --- |
| Passive-Aggressive Classifier | 0.85 |
| SGD Classifier with Log Loss | 0.91 |
| Perceptron | 0.89 |
| Linear SVM (One-vs-Rest) | 0.91 |
| Linear SVM (LibLinear Implementation) | 0.92 |
| L1-Regularized Logistic Regression | 0.92 |
| L2-Regularized Logistic Regression | 0.91 |
| One-vs-All Linear Regression | 0.91 |
| Multinomial Logistic Regression | 0.92 |
| Ridge Classifier | 0.91 |
| Linear Discriminant Analysis (LDA) | 0.94 |

We chose logistic regression with a One-vs-All (OvA) strategy to achieve the main goals of our research. This choice is motivated by the importance of being able to clearly distinguish each brain state from all others during classification. Technically, in the OvA approach, multiclass classification is performed by training *s* independent binary classifiers, one for each brain state class *c* = {1, …, *s*} . Each such classifier is trained to distinguish one brain state class *c* from all others by computing a corresponding weight vector 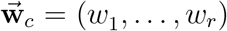. This vector represents the contribution of each feature — corresponding to the mean activity of a specific brain region — to the identification of brain state *c*.

Thus, for each of the *s* classifiers, the weights *w*_*i*_ reflect the importance of the corresponding features for a specific brain state class: the larger absolute value |*w*_*i*_|, the stronger the influence of the *i*-th brain region on the prediction of membership in brain state class *c*. This directly enables us to analyse the contribution of individual brain regions to the discrimination of each specific state from all others, which is a key requirement for neurobiological interpretation.

For a given input vector of mean activity values 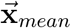 , the probability of belonging to brain state *c* is calculated using the function:

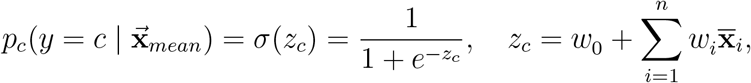

where:

- *w*_*i*_ is the weight of the *i*-th brain region for brain state *c*,
- *w*_0_ is the bias term (intercept) for brain state *c*.

As previously mentioned, each classifier produces a set of posterior probabilities 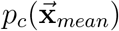, where 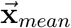 denotes the feature vector and *c* ∈ {1, 2, … , *s*} represents the possible brain state class labels. The final predicted brain state class *c* is determined by selecting the class with the highest posterior probability, i.e., the class deemed most probable given the observed features:

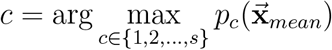

The classification results for each brain state class are summarised in Table 3. For each brain state class, we report both the class-specific accuracy and the actual number of test samples. The total dataset comprised 8134 measurements. We split it into 80% for training and 20% for testing, resulting in a test set size of 1638 measurements. To prevent data leakage across subjects, the train/test split was performed at the subject level using a group-based splitting strategy, ensuring that all measurements from a given subject were assigned exclusively to either the training or the test set. This procedure resulted in exactly 117 subjects per class being allocated according to the same subject-level partitioning scheme, thereby preserving class balance across splits.The model’s Accuracy of 0.9 (from Table 2) indicates it correctly predicted 90%. All analyses were conducted in Python using `scikit-learn`. The classifier was trained with the following hyperparameters: maximum number of iterations (`max_iter = 1000`), default regularization parameter (`C = 1.0`) and `L2` penalty. The data were standardized using `StandardScaler` prior to model fitting. Train/test splits were performed with `random_state = 42` to ensure reproducibility.

**Table 3:**
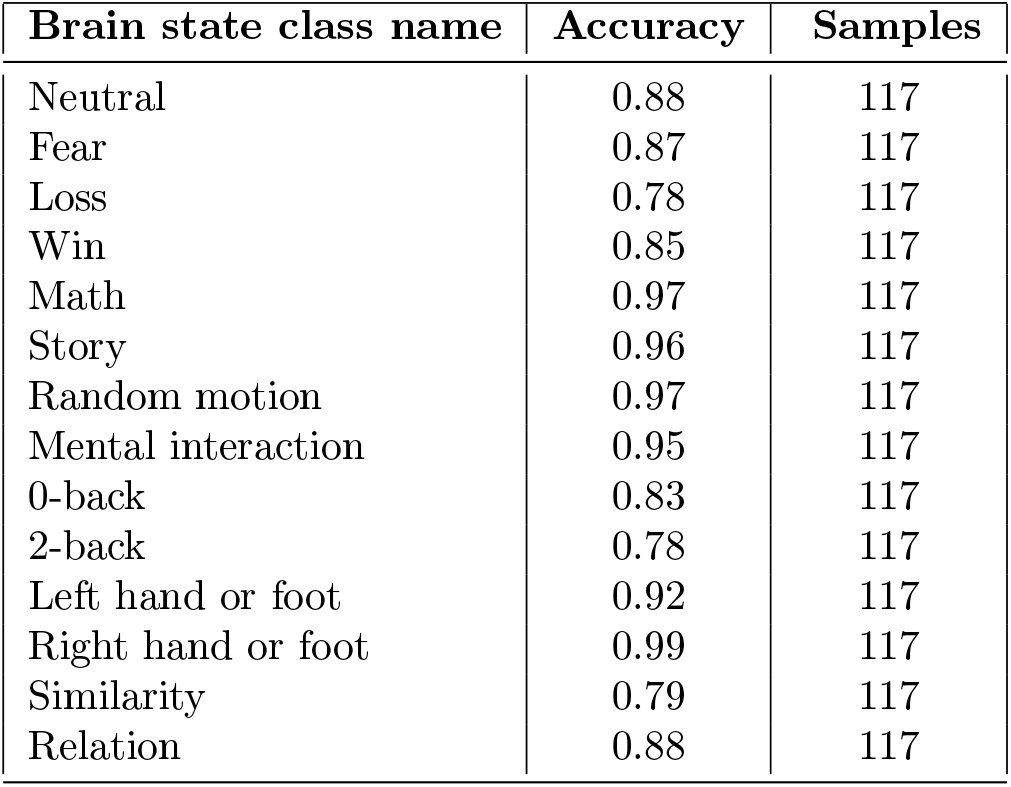
Classification Report with OvA.

To obtain a more reliable estimate of model generalization performance beyond a single train/test partition, we additionally performed 5-fold group cross-validation at the subject level. In each fold, subjects were split into training and validation sets without overlap, and data standardization was performed within the training fold only. The classification accuracies across folds were 0.8962, 0.8904, 0.8824, 0.8910, and 0.8842, yielding a mean accuracy of 0.8888 with a standard deviation of 0.0056 across folds. The low variability across folds indicates stable performance across different subject-level splits and supports the model’s generalization capability.

To provide statistical validation of the reported performance, we conducted two complementary analyses for the One-vs-Rest (OvR) logistic regression classifier. First, we applied bootstrap resampling of the test set with replacement (1000 iterations). For each bootstrap sample, classification accuracy was recomputed, yielding an empirical distribution of accuracies. From this distribution we derived a 95% confidence interval, which placed the observed accuracy of 0.921 within the range [0.907, 0.933]. This indicates that the model’s performance is robust and not attributable to sampling variability.

To contextualize the observed classification performance relative to chance level, we estimated the baseline accuracy using two complementary approaches.

First, we computed the empirical null distribution via a permutation test. The test set labels were randomly permuted 1000 times while keeping model predictions fixed, and classification accuracy was recalculated for each permutation. The resulting null distribution had a mean accuracy of 0.0717 (standard deviation is 0.0062), which corresponds to the expected chance-level performance for a 14-class classification problem. The observed accuracy of 0.9188 lies far outside this null distribution, yielding a permutation test *p*-value < 0.001.

Second, we evaluated a DummyClassifier (most frequent class strategy), which achieved an accuracy of 0.0714, consistent with the theoretical random baseline (1/14 ≈ 0.0714).

Both baseline estimates are substantially lower than the observed model accuracy, confirming that the classifier performs far above chance level and that the reported performance cannot be attributed to random label assignment.

We highlight the high precision of the states — *left hand or foot, and right hand or foot, math, story, mental interaction, random motion* — which correspond to experiments in **Motor Task** for mapping brain motor regions, **Language Processing** for studying brain areas involved in language comprehension and arithmetic processing, and **Social Cognition** for identifying neural mechanisms of social interaction and intention recognition.

### 2.5 Significant Brain Regions Identification

To identify the most significant brain regions which contribute the most to the classification of different brain states, we employ the following approach.

#### 2.5.1 Analysis of Model Accuracy

For each brain state class *c* = {1, …, *s*}, we rank the features – averaged brain regions signals 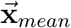, according to the absolute values of their corresponding weights in 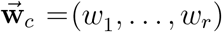. This procedure established a distinct feature importance ranking for each brain state class.

To analyse the dependence of model accuracy on the number of used features, we applied the following approach: we iteratively removed features, starting from the least significant ones. At each iteration, we retrained the model on the reduced feature set, re-ranked the remaining features by their updated absolute weights, and evaluated per-class performance using the True Positive Rate (TPR or Recall). We chose TPR because it directly measures how feature reduction impacts the model’s ability to detect all instances of each class. As TPR quantifies the proportion of target examples correctly identified, it is critical for our task where missing true positives is unacceptable. The TPR values (averaged across classes) were recorded and visualized on the Fig. 1, with the *X*-axis showing the number of remaining features and the *Y* -axis showing the corresponding accuracy.

**Fig. 1:**
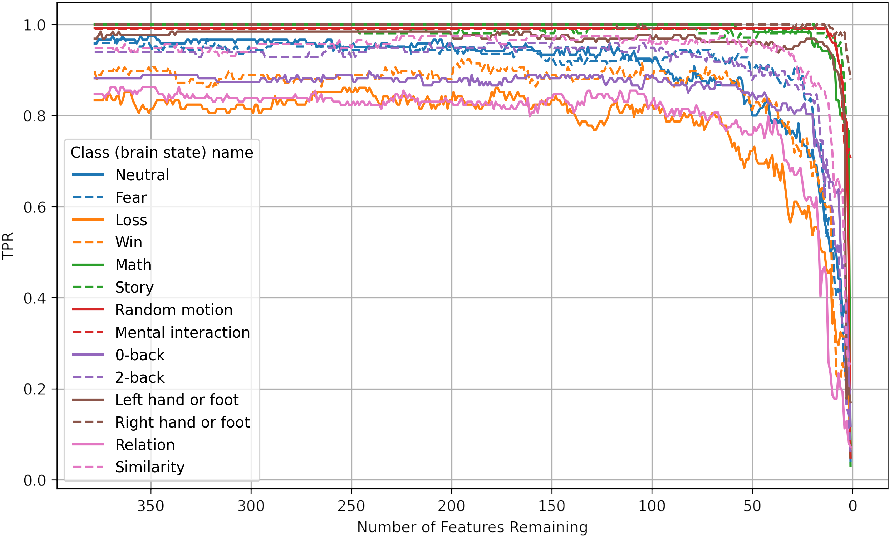
Accuracy plot depending on the number of features

This graph clearly demonstrates that feature selection can be performed without significant loss of accuracy up to a certain limit. However, as the number of features is further reduced, the model begins to lose its ability to classify adequately, highlighting the importance of selecting the right number of features to maintain a balance between model simplicity and its quality.

#### 2.5.2 Significant Brain Regions Selection

We begin with the complete set of significant brain regions and iteratively remove features in descending order of importance, as determined by absolute weight values. After each removal step, we: (1) evaluate the model’s true positive rate (TPR), and (2) re-rank the remaining features based on their updated weights in the refined model. If the TPR drops by more than a specified threshold (we tested for 5% and 10%) relative to the maximum value observed prior to feature elimination, the last removed feature is restored and the procedure terminates. This criterion ensures that feature elimination continues only while the TPR remains within acceptable limits. This method allows us to retain key brain regions that significantly impact the model’s TPR while eliminating less significant features, minimizing the loss in classification quality.

As a result, we obtained an informative set

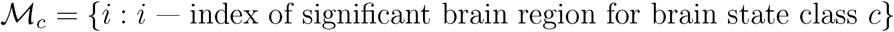

of significant brain regions for each brain state class *c* = {1, …, *s*}, and these sets will be further analysed.

Fig. 2 presents the distribution of significant brain regions set sizes across different classes at accuracy reduction thresholds of 5% and 10%. The results demonstrate substantial inter-class variability in the number of brain regions required for effective classification. Classes exhibiting high classification accuracy consistently show compact sets of significant brain regions at both thresholds, indicating that these brain states are characterized by specific, localized neural activity patterns that facilitate robust discrimination.

**Fig. 2:**
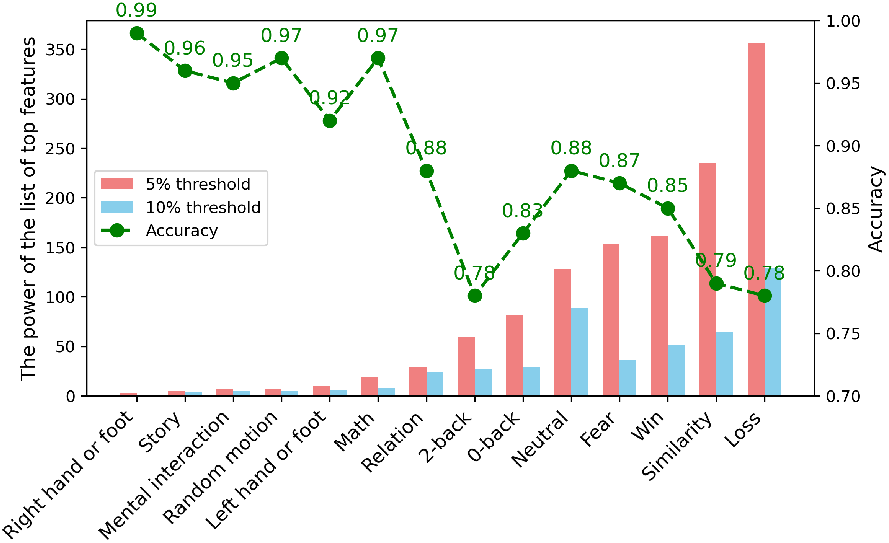
The graphs of cardinality of the top features sets

Conversely, states with lower classification accuracy require substantially larger feature brain regions for characterization. This pattern holds true at both examined thresholds, suggesting that these brain states lack clearly identifiable focal signatures. Instead, their neural representations appear to involve distributed activity across multiple cortical areas, necessitating integration of information from broader neural networks for accurate classification.

#### 2.5.3 Unique Sets of Significant Brain Regions

Our analysis uses feature sets generated at a 5% accuracy drop threshold. To quantify the similarity between the sets of significant brain regions corresponding to different brain state classes, we computed the Jaccard coefficient for each pair of classes *i* and *j*. The Jaccard coefficient is computed as the ratio of the intersection of sets to the union of sets:

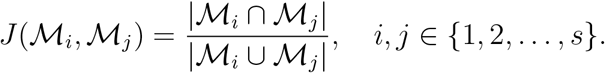

The Jaccard coefficient ranges between [0, 1], where 1 indicates that the sets are identical, 0 indicates that the sets have no common elements.

By pairwise comparing the sets of significant brain regions, we construct a Jaccard coefficient matrix:

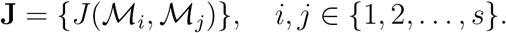

The Jaccard coefficient matrix **J** is a symmetric matrix of size *s* × *s*. This matrix allows us to assess the degree of overlap between significant brain regions across different brain state classes.

Furthermore, Fig. 3 shows that the sets of significant brain regions ℳ_*i*_, corresponding to brain states with high classification accuracy, hardly overlap. This indicates that for each brain state, the model has identified a unique set of regions whose mean activity is most significantly associated with that state. However, there are also sets whose elements are similar in composition. This occurs when these sets approach the comprehensive coverage of the HCP MMP atlas (*r* = 379 regions), resulting in reduced specificity of significant region identification. Under these conditions, the distinctive contributions of individual regions become less discernible, complicating the interpretation of their functional roles in state classification. Therefore, we focus our sequential analysis on six conditions — *math, story, random motion, mental interaction, left hand of foot, right hand or foot*.

**Fig. 3:**
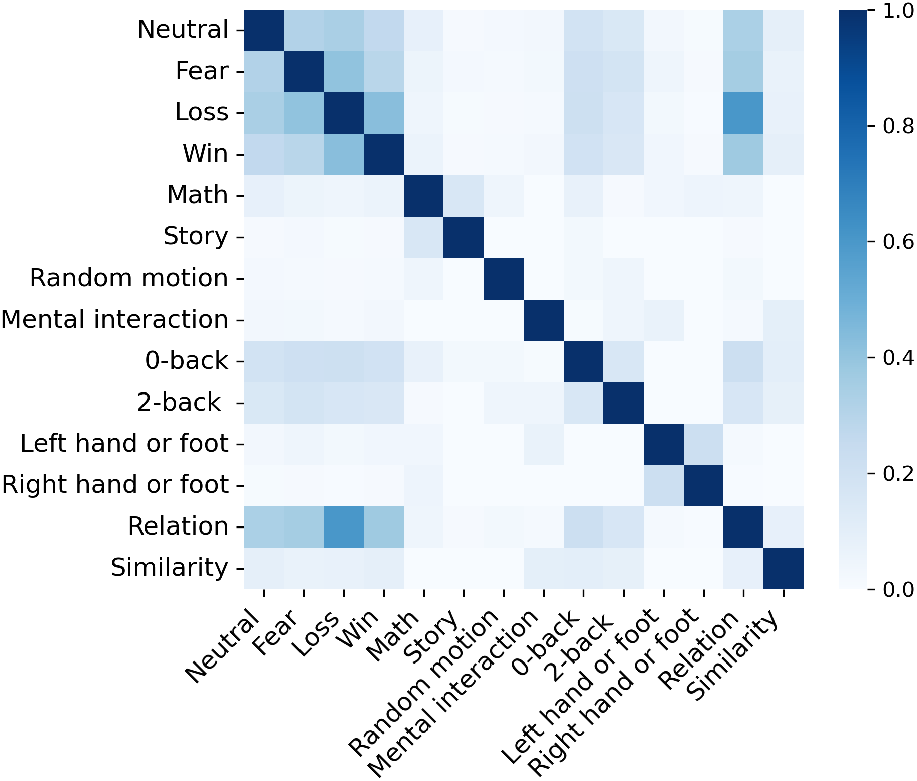
The Jaccard coefficient matrix **J** heatmap

The proposed approach not only allows us to identify the most significant brain regions for each condition but also reveals the relationship between the number of significant brain regions and classification accuracy. This highlights the importance of analysing the spatial structure of brain activation to understand the neurobiological mechanisms underlying different states.

The identified significant brain regions, their corresponding weights, the correlations between them, as well as their neurobiological interpretation are presented in Section 3.1.

### 2.6 Analysis of Temporal Dynamics

For the classes with high classification accuracy — *Math, Story, Random Motion, Mental Interaction, Left Hand or Foot, and Right Hand or Foot* — our sequential analysis focuses on these six brain conditions. We focus on such classes because the small number of significant brain regions indicates the presence of clearly defined and localized patterns of brain activation that play a key role in the formation of these states.

One of the main tasks of our analysis is to verify the significance of the temporal structure in identifying functional connections between brain regions. We worked with fMRI data, where each significant region is represented by a time series. By studying the correlations between these time series, we aimed to determine whether temporal dynamics significantly contributed to the formation of functional connections or whether these connections can be identified based on averaged or static characteristics of activity.

To analyse in more depth the importance of temporal structure in functional connections between significant brain regions, we perform the following steps, as described in sections 2.6.1, 2.6.2.

#### 2.6.1 Temporal Correlation Analysis

For each activity matrix **X** corresponding to brain state class *c*, we perform the following steps.

1. We extract the time series 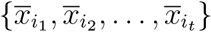 for significant brain regions, where the region indices belong to the set

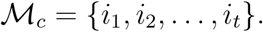
2. We construct the correlation matrix 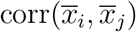, where *i, j* ∈ ℳ_*c*_ are the indices of the selected significant brain regions of the extracted time series, where each element represents the Pearson correlation coefficient between two time series.

Constructing correlation matrices of time series allows us to study the relationship between the activities of different brain regions that have been identified as significant for each state *c*. Pearson correlation coefficients show how closely the activity of one region changes relative to another within a single observation. High correlation values may indicate synchronous activation of regions, which could suggest their functional relationship in the context of the given brain state.

For each pair (*i, j*) of significant brain region indices *i, j* ∈ ℳ_*c*_ we construct

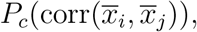

the distributions of the Pearson correlation coefficients between time series of significant brain regions *i, j* ∈ ℳ_*c*_ for all activity matrices of class *c*.

Analogously, we define 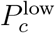 as the distributions of correlations for the low-ranked brain regions, selected in equal number per class for comparability.

Analysing the distributions allows us to assess the stability and nature of interactions between pairs of brain regions within a single state. For example, if the distribution of correlation coefficients between two regions shows a consistently high value, this may indicate their systematic co-activation. In contrast, a wide or sparse distribution may suggest variability in the connection between these regions.

We analysed the correlations between the time series of highly informative brain regions, selected based on their contribution to the classification model. We compared the distributions of temporal correlations between brain regions identified as most significant by the classifier (*P*_*c*_) and those ranked lowest 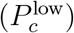 As shown in Fig. 4, the two distributions differ clearly: correlations among top-ranked regions are shifted toward higher values, whereas low-ranked regions exhibit substantially weaker correlations.

**Fig. 4:**
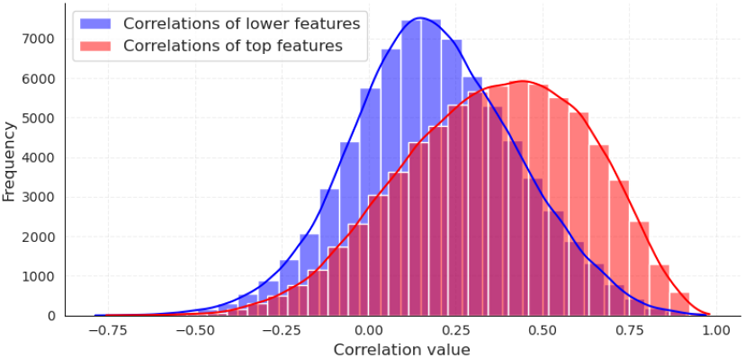
Distributions of time series correlation of top-ranked and low-ranked regions

To quantitatively assess the differences between the correlation distributions of top-ranked and low-ranked brain regions, we reformulated the problem as a binary classification task. In this setup, distributions *P*_*c*_ for top-ranked regions and for 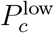 low-ranked regions were assigned to two separate classes. We then constructed a receiver operating characteristic (ROC) curve by evaluating the trade-off between the true positive rate (TPR) and false positive rate (FPR) across different correlation thresholds. This approach allows us to quantify how well the correlation structure of top-ranked regions can be distinguished from that of low-ranked regions, thereby providing a robust measure of separability between the two distributions. The resulting ROC analysis, along with the descriptive statistics of the distributions, is presented in Section 3.2.

#### 2.6.2 Analysis of Temporal Structure Significance

In the original data, the time series of the significant brain regions 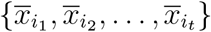 reflect the dynamic activity of specific brain regions, and their temporal dependencies play an important role in the formation of functional patterns and interactions between regions. In each extracted subset of the time series 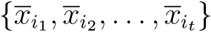 , we shuffled all time series to create a control dataset where the temporal structure of brain region activity is completely disrupted, and the original dependencies within the time series are eliminated.

To denote the shuffled values of the time series we introduce the following notation:

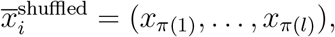

where *π* : {1, 2, … , *l*}→{1, 2, … , *l*} is a random permutation of the time point indices.

As shown in Fig. 5, the permutation *π* reorders the elements of the series of length *l* = 4.

**Fig. 5:**
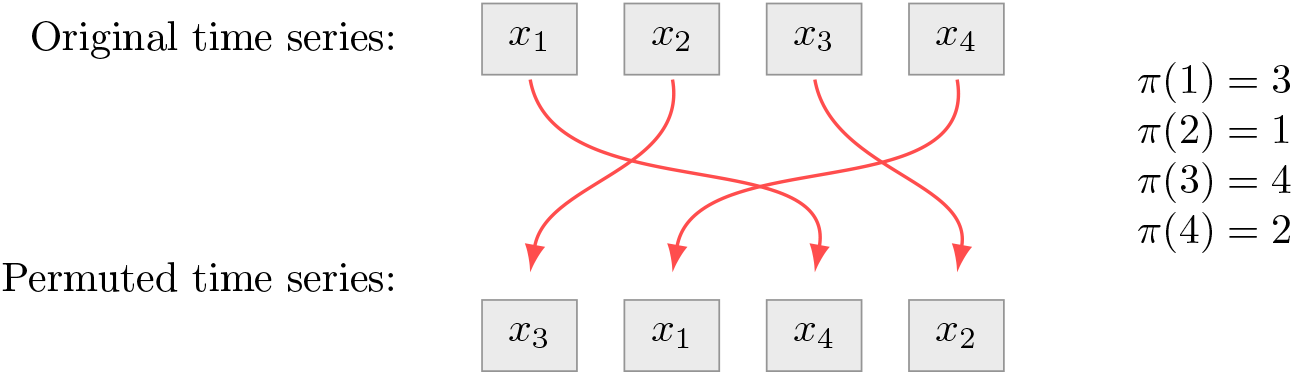
Example of the permutation *π* for a time series of length *l* = 4.

In the shuffled series there is no temporal structure, which eliminates the influence of dynamic coordination between brain regions.

Similarly to the previous section, we construct

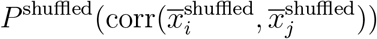

— the distribution of the Pearson correlation coefficients between the shuffled time series of the significant brain regions *i, j*, ∈ ℳ_*c*_ for all activity matrices of class *c*.

We compared the distributions of the original and shuffled time series to determine which properties of temporal activity are truly critical for the formation of functional patterns associated with brain states. To analyse the differences between distributions and assess the influence of temporal structure on correlation coefficients, we perform the following analysis for each significant brain regions pair (*i, j*) where *i, j*, ∈ ℳ_*c*_. We conducted a Kolmogorov-Smirnov (KS) test to compare: the distribution of Pearson correlation coefficients from the original time series of the significant brain regions, versus the distribution from the shuffled time series. Formally, the KS test examines the hypothesis that the two distributions originate from the same underlying population.

The KS test statistic is defined as:

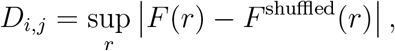

where:

- 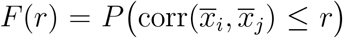 is the correlation distribution function of the original time series,
- 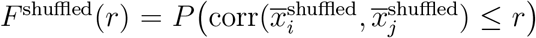 is the correlation distribution function of the shuffled time series.

Based on the value of the test statistic *D*_*i,j*_ and the corresponding *p*-value, we can assess the significance of the differences between the distributions. We set the significance level at *α* = 0.03. If the *p*-value is less than *α*, the null hypothesis of equal distributions is rejected, indicating significant differences in the correlation structure between the original and shuffled time series for a given pair of features *i* and *j* in class *c*. Thus, for each pair of features *i* and *j* within class *c*, we obtain the KS statistic *D*_*i,j*_ and the corresponding *p*-value, allowing us to evaluate the significance of differences in the correlation structure.

## 3 Results

### 3.1 Significant Brain Regions

Our model identified unique sets of brain regions whose activity is most strongly associated with the six conditions with high classification accuracy.

The HCP indices of the brain regions significant for classification, along with their anatomical labels are provided in Table 4. The results are accompanied by detailed visualizations showing both model weights and mean activation values across these critical regions (Fig. 6). The absolute value of the weight reflects the strength of the feature’s influence — the higher it is, the more significant the region’s contribution to determining the state. The sign of the weight indicates the nature of this influence: a positive value means that increased activity in the region raises the probability of assigning the object to the given state, while a negative value means that increased activity, conversely, reduces it. By analysing the signs of the model weights for the selected regions, we can identify activators and inhibitors of the processes associated with each brain state.

**Table 4:**
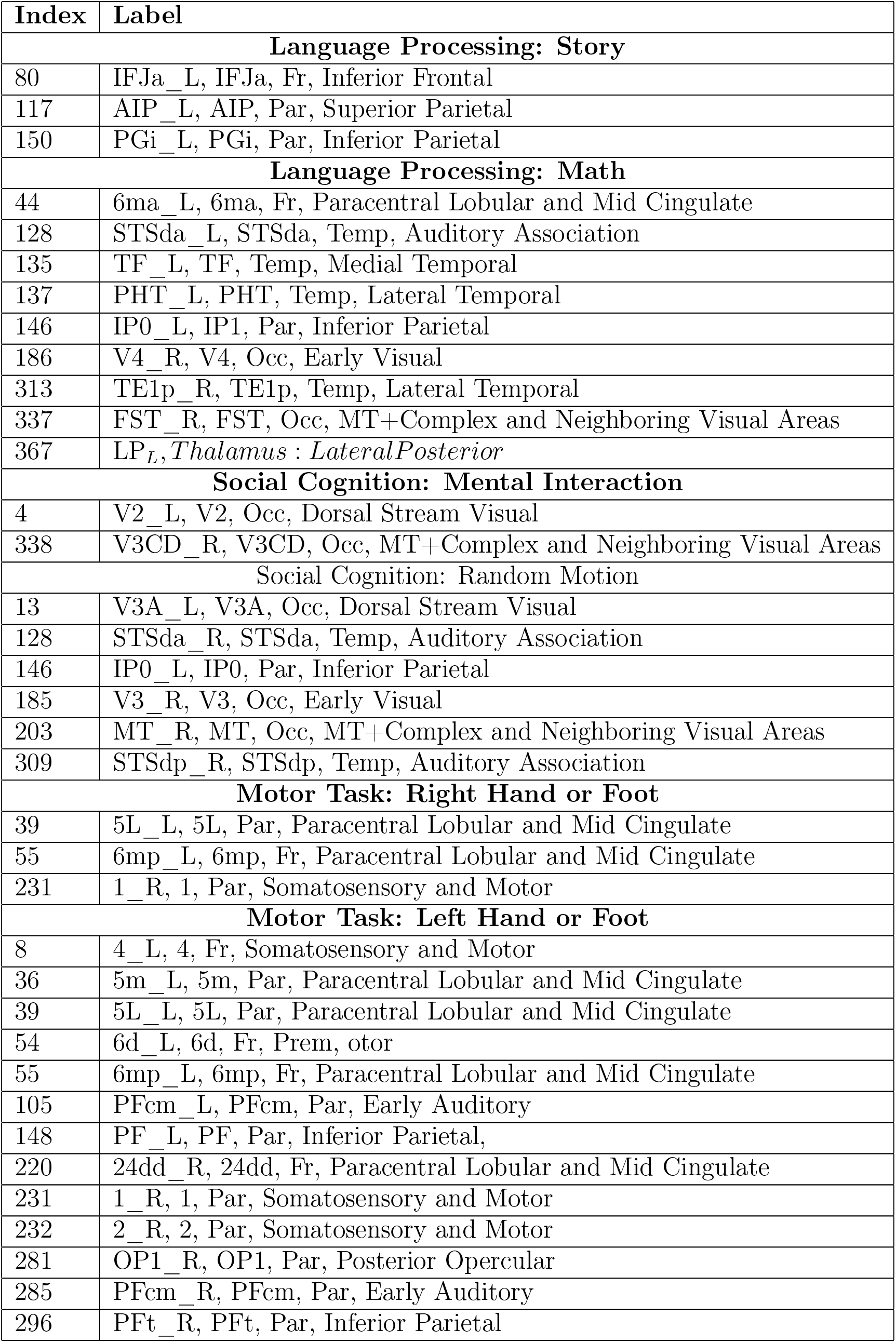
The HCP indices and anatomical labels of significant brain regions.

**Fig. 6:**
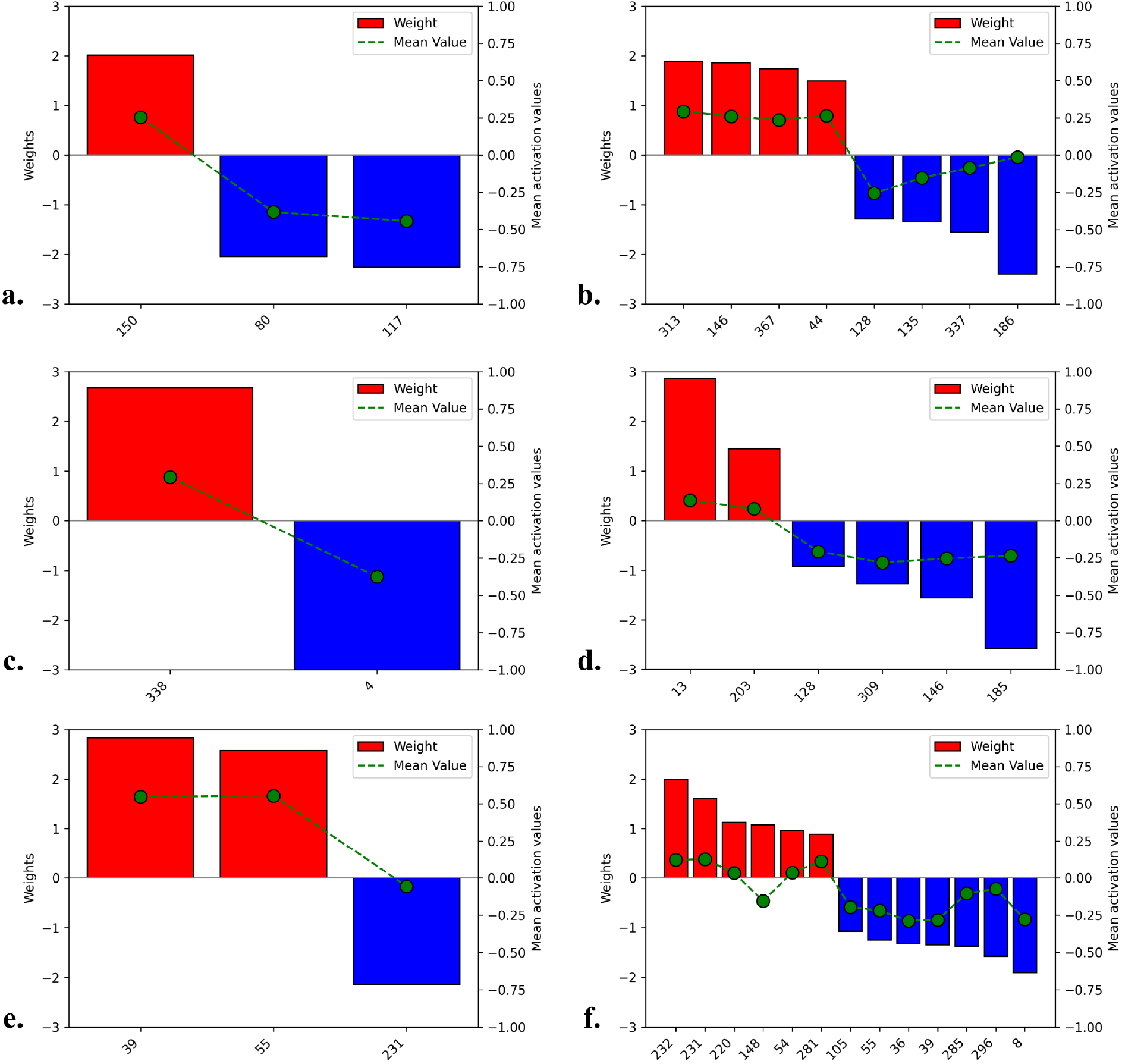
Significant brain regions weights and mean activation values for the six task-induced brain states with high classification accuracy. **a**. Language Processing: Story **b**. Language Processing: Math **c**. Social Cognition: Mental Interaction **d**. Social Cognition: Random Motion **e**. Motor Task: Right Hand or Foot **f**. Motor Task: Left Hand or Foot

Additionally, for each state, we included a heatmap showing correlations between key regions (Fig. 7). The color scale visualizes the strength and direction of the connections: red indicates a strong positive correlation, white represents no correlation, and black signifies a strong negative correlation. This allows for a clear assessment of the degree of interaction between different brain areas within the context of the studied states.

**Fig. 7:**
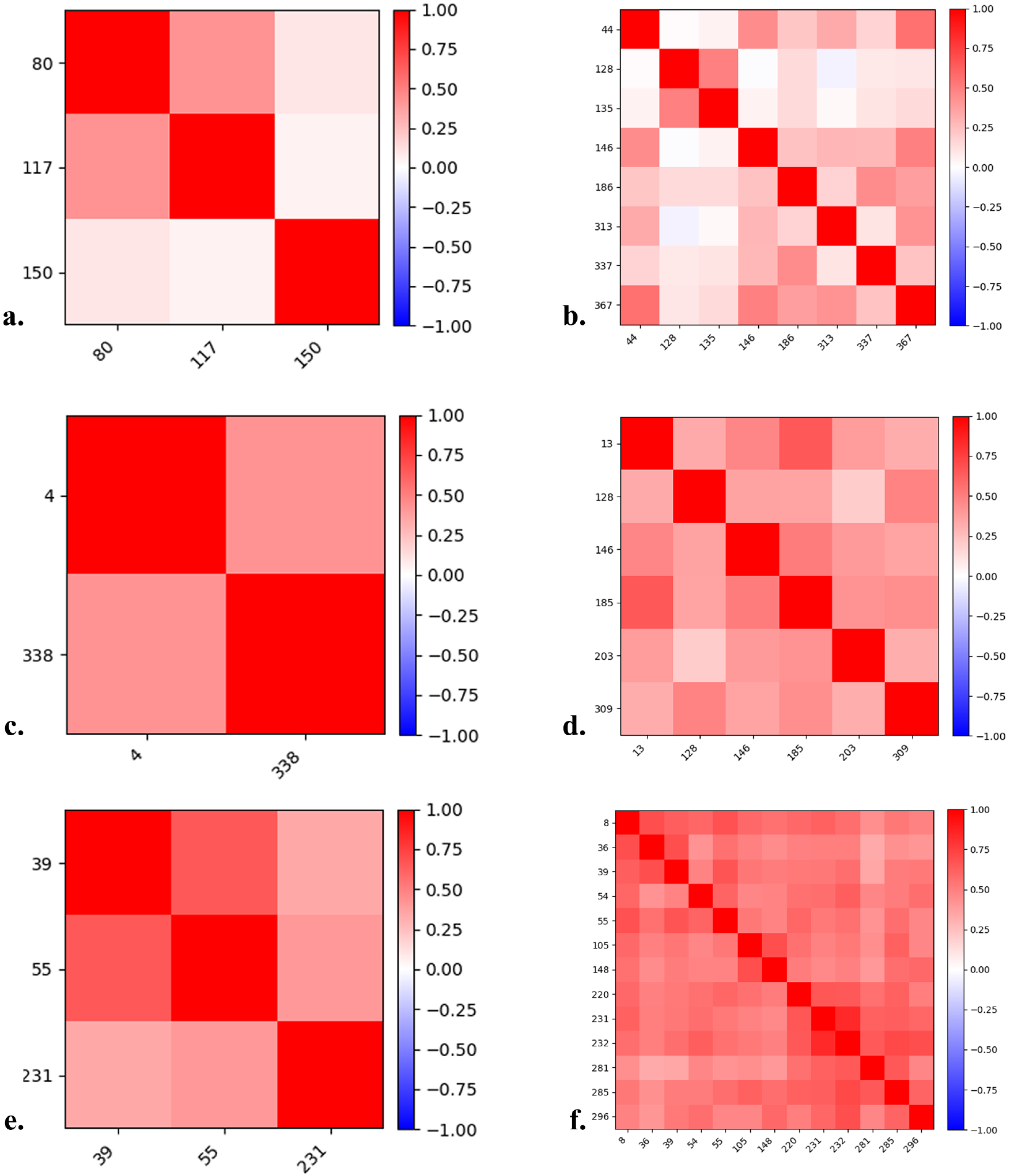
Heatmaps of Pearson correlations between regional mean activations of significant brain regions for the six task-induced brain states with high classification accuracy. **a**. Language Processing: Story **b**. Language Processing: Math **c**. Social Cognition: Mental Interaction **d**. Social Cognition: Random Motion **e**. Motor Task: Right Hand or Foot **f**. Motor Task: Left Hand or Foot

#### Language Processing: Story

Fig. 6a shows the weights and mean activation values of key brain regions most significant for classifying the Story state. Region 150 (left PGi) functions as an activator, while region 80 (left IFJa) and region 117 (left AIP) exhibit an inhibitory effect — their activity reduces the likelihood of classifying the current state as Story.

The heatmap in Fig. 7a demonstrates correlations between the three significant brain regions. The most pronounced positive correlation (shown in red) is observed between the left AIP and IFJa.

#### Language Processing: Math

Fig. 6b shows the weights and mean activation values of the key brain regions most predictive of the Math brain state.

Region 313 (right TE1p) exhibits the highest positive weight, highlighting its primary role as an activator during arithmetic problem solving. Stronger — but still somewhat lower — activating effects are also seen in region 146 (left IP0), region 367 (left LP thalamus), and region 44 (left 6ma).

In contrast, region 186 (right V4) displays the strongest inhibitory effect: its activation decreases the probability of classifying the state as Math. Additional inhibitors include region 128 (left STSda), region 135 (left TF) region 337 (right FST). Mean BOLD-signal values confirm this pattern: activators have positive mean activations, whereas inhibitors have negative means (the lowest being in regions 186, 128, and 135).

Fig. 7b presents the pairwise correlations among these key regions. The strongest positive correlation is between the left 6ma and left LP thalamus, indicating tight functional coupling during numerical information processing. Strong correlations are also observed between the left 6ma and left IP0, as well as between the left IP0l and LP thalamus, underscoring their synchronized involvement in the visuospatial and numerical components of arithmetic computation.

#### Social Cognition: Mental Interaction

Fig. 6c shows the weights and mean activation values of the key brain regions most predictive of the Mental Interaction state.

Region 338 (right V3CD) exhibits a strong positive weight, marking it as the primary activator: increased activation here is tightly linked to perceiving intentional interaction. In contrast, Region 4 (left V2) carries a strong negative weight, acting as an inhibitor—higher activity in this area decreases the likelihood of classifying the trial as Mental Interaction. Mean BOLD-signal values mirror these roles: region 338 shows a high positive mean activation, whereas mean activation of V2 is negative.

Fig. 7c presents the pairwise correlations between these two regions. A clear positive correlation between the left V2 and the right V3CD indicates their synchronized engagement when observers infer purposeful interactions among moving shapes.

#### Social Cognition: Random Motion

Fig. 6d shows the weights and mean activation values of the key brain regions most predictive of the Random Motion state, in which shapes moved without any apparent interaction.

Region 13 (left V3A) carries the highest positive weight, marking it as the principal activator when participants perceive purely stochastic motion. A secondary activator is region 203 (right MT). All other regions act as inhibitors: the mildest inhibitory effect appears in region 128 (right STSda), followed by region 309 (right STSdp) and region 146 (left IP0), with region 185 (right V3) showing the strongest inhibitory weight. Mean BOLD-signal values mirror these roles: activators (left V3A and right MT) have positive mean activations, whereas inhibitors (right STSda, right STSdp, left IP0, right V3) all exhibit negative means, deepest in region right V3.

Fig. 7d presents the pairwise correlations among these six regions. The strongest positive correlation is between the right MT and right STSdp, while the weakest occurs between the right STSda and right MT. Overall, all off-diagonal correlations are positive—ranging from moderate to strong — indicating a broadly co-activated visual–association network even under random motion.

#### Motor Task: Right Hand or Foot

Fig. 6e shows the weights and mean activation values of the key brain regions most predictive of the Right Hand or Foot motor state.

Region 39 (left 5L) carries the highest positive weight, marking it as the primary activator during contralateral hand or foot movements. Region 55 (left 6mp) follows closely as a secondary activator. In contrast, region 231 (right area 1) exhibits a strong inhibitory weight, indicating that its activity decreases the probability of classifying the state as Right Hand or Foot. Mean BOLD-signal values align with these roles: left 5L and left 6mp show positive mean activations, whereas the right area 1 has a negative mean.

Fig. 7e presents the pairwise correlations among these three regions. The strongest positive correlation occurs between the left 5L and left 6mp, reflecting their joint engagement in motor planning and execution with a dominant right side. The weakest correlation is between the left 5L and right area 1, though all off-diagonal values are positive and moderate, indicating overall co-activation across paracentral motor and somatosensory areas during right-side limb movements.

#### Motor Task: Left Hand or Foot

Fig. 6f shows the weights and mean activation values of the key brain regions most predictive of the Left Hand or Foot motor state.

Region 232 (right area 2) demonstrates the highest positive weight, identifying it as the principal activator for left-side movements. Close behind are region 231 (right area 1) and region 220 (right 24dd), followed by region 105 (left PFcm), region 148 (left PF), region 54 (left 6d), and region 281 (right OP1), all of which also contribute positively to classifying the “Left Hand or Foot” state.

In contrast, region 8 (left area 4) carries the strongest negative weight, acting as the main inhibitor; additional inhibitory effects are seen in region 285 (right PFcm), region 296 (right PFt), region 39 (left 5L), region 36 (left 5m), and region 55 (left 6mp). Mean BOLD-signal values mirror these roles: all activators show positive average activations, whereas inhibitors exhibit negative means (deepest in the left area 4).

Fig. 7f presents the pairwise correlations among these regions. The strongest positive correlation is between the right area 2 and right area 1, reflecting their joint recruitment during left-side motor control. The weakest (yet still positive) correlation occurs between the left 5L and right PFcm. Overall, all off-diagonal correlations are positive and range from moderate to strong, highlighting a cohesive network engaged by left-side limb movements.

### 3.2 Analysis of Temporal Dynamics

Finally, for each brain state class, we performed the steps described in Sections 2.6.1, 2.6.2.

As described in Section 2.6.1, we constructed the empirical distributions of temporal correlations between the time series of top-ranked and low-ranked brain regions. For the low-ranked regions, the correlation distribution had a mean of 0.188, median 0.181, and standard deviation 0.237, with the central 50% of values spanning [0.030, 0.347]. By contrast, the top-ranked regions showed a higher mean correlation of 0.360, median 0.376, and standard deviation 0.264, with the central 50% spanning [0.176, 0.562].

To quantitatively assess the difference between these distributions, we treated them as representing two distinct classes: class 0 for the top-ranked regions and class 1 for the low-ranked regions. We then constructed a receiver operating characteristic (ROC) curve by comparing TPR and FPR across varying thresholds. The resulting AUC = 0.7 confirms that the correlation distribution of top-ranked regions offers stronger class separation.

These results indicate that brain regions most informative for classification are also more strongly correlated in their temporal dynamics compared to low-ranked regions, reinforcing the interpretation that the discriminative states emerge from coordinated activity patterns in specific neural circuits.

Section 2.6.2 outlined our approach to testing the influence of temporal structure by comparing original and shuffled time series. The Kolmogorov-Smirnov (KS) test results demonstrate that for nearly all feature pairs (*i, j*) in each class, the *p*-values were significantly below the chosen significance threshold of *α* = 0.03. This confirms that the distributions of correlation coefficients from original time series are statistically distinct from those derived from shuffled time series. However, for the state *Motor Task: Left Hand or Foot*, two feature pairs did not pass the significance threshold: regions (105, 232) with a *p*-value of 0.06125, and regions (281, 296) with a *p*-value of 0.09603, both greater than 0.03.

## 4 Discussion

The present findings are based exclusively on data from the Human Connectome Project (HCP). While this dataset provides high-quality, standardized acquisitions, the generalizability of the reported results to data acquired with different scanners, acquisition protocols, or to other populations remains to be established. Future studies are needed to validate the robustness of the proposed approach across independent datasets and more diverse cohorts.

A key focus of our research was to demonstrate that classical, somewhat mundane ML algorithms can successfully classify complex task-based fMRI data. By applying these algorithms, we identified significant brain regions whose activity was strongly correlated with a specific condition. We found unique sets of regions for each task-induced brain state, with minimal overlap between classes (confirmed by the Jaccard coefficient analysis). The ML approach showed us once again that each task-induced brain state is characterized by a distinctive activity pattern arising from specific neural network dynamics.

Our ML classification approach had proven itself to be technically robust and effective. However, the results obtained had to be evaluated against established neuroscientific knowledge. In the following, we provide an interpretation of the significant brain regions obtained for the six brain states.

### Language Processing: Story

The Story task consisted of listening to a fable-like story and answering a forced-choice question about the topic of the story. We identified three key brain regions – all in the left hemisphere – that are most significant for classifying the Story state. The contribution of the left PGi was positive, meaning that increased activity in this region promotes classification into the Story state. The PGi is the most anterior and inferior part of the inferior parietal cortex. Its functional connectivity implicates PGi in the ventral stream of visual/auditory information processing (the “what” stream). Moreover, its strong linkages to temporoparieto-occipital junction areas — which are centrally involved in theory of mind processing — suggest a critical role in deciphering social cues and inferring others’ mental states. Simultaneously, PGi bridges these multimodal representations with memory systems via posterior cingulate cortex connectivity, creating a neural interface where socially relevant sensory inputs interact with episodic and semantic memory frameworks. Hence, the PGi is implicated in multiple, complex cognitive processes – such as semantic integration, social cognition, and theory of mind – that are associated with the default mode network (DMN) functioning and targeted by the Story task (Jääskeläinen et al., 2020).

However, the activation of the left IFJa and AIP – located in the lateral prefrontal cortex and intraparietal sulcus respectively – reduced the likelihood of classification into the Story state. The IFJa is a key node in the central executive network, associated with task-set reconfiguration and attentional shifts (Sundermann & Pfleiderer, 2012), while the AIP supports sensorimotor integration and action planning (Bisley et al., 2011). The strong positive correlation between AIP and IFJa (Fig. 7a) may suggest that these regions operate as a coordinated inhibitory circuit, dynamically suppressing DMN activity during goal-directed tasks.

#### Language Processing: Math

The math task required participants to listen to addition and subtraction problems and then respond by choosing the correct answer from two options. Four regions in the left hemisphere, including the TE1p (a part of the auditory cortex), IP0 (a region in the inferior parietal cortex), the lateral posterior (LP) nucleus of the thalamus, and area 6ma (a premotor region) — showed significant activity patterns that helped distinguish when participants were engaged in the math task compared to other states. These findings align well with existing literature about neural correlates of mental arithmetic, which consistently indicates a specialized but distributed network of cortical and subcortical brain regions (Rosenberg-Lee et al., 2011; Salillas et al., 2021). Even the premotor areas (6ma), seemingly unrelated to mathematical reasoning, are found to be crucial for mental computation through metaphorical motion and structure mapping processes (Andres et al., 2011). Also, this particular task was rather complex. The numerical expressions were presented in an unusual auditory form and the calculations had to be done in mind. Hence, the involvement of the auditory areas (TE1p).

In contrast, the right V4 and left TF (a region within the parahippocampal gyrus), along with the right FST and left STSda (both situated in the superior temporal sulcus), exhibited an inhibitory effect, whereby increased activation in these areas decreased the likelihood of classifying the brain state as Math. This pattern suggests that during mental arithmetic, which demands focused abstract and executive processing, regions involved in detailed visual perception (V4 and FST) and higher-order social or memory-related processing (STSda and TF) are downregulated. The suppression of V4 and FST likely reflects a functional disengagement from sensory-driven visual analysis, allowing cognitive resources to be reallocated toward numerical manipulation and calculation. Similarly, decreased activity in STSda and TF may indicate reduced engagement of social cognition and episodic memory processes, which are less relevant during focused mathematical computation. These findings align with prior evidence that mathematical tasks preferentially activate frontoparietal networks while attenuating activity in sensory and default-mode regions to optimize task performance (Ylinen et al., 2024).

#### Social Cognition: Mental Interaction

During this task, participants were presented with moving geometrical shapes and had to judge whether simple shapes were intentionally interacting or moving at random. Here, the increased activation of the right V3CD was related to perceiving intentional interaction. In contrast, a higher activation of the left V2 decreased the likelihood of classifying the trial as Mental Interaction.

The V3CD, as part of the dorsal visual stream (the “where” stream), is involved in processing complex motion and shape information, contributing to the perception of moving objects. Its increased activation during judgments of intentional interaction suggests that this region supports the integration of motion cues necessary for conscious visual perception and motion analysis (Pitzalis et al., 2010).

Although the V2 is also involved in global motion processing (among other functions), its inhibitory effect on classification implies that when early-stage visual processing dominates, the perception of intentional interaction is less likely.

#### Social Cognition: Random Motion

During the random motion runs, the participants were presented with similar visual stimuli, but without any interaction between the objects. Surprisingly, this state involved a much larger network of key brain regions, both with positive and negative weights.

Here, only the left V3A and the right MT showed significant activation patterns that helped to distinguish the Random Motion state. All other regions – the right STSda and STSdp (subparts of the superior temporal sulcus), together with the left IP0 and the right V3 – showed a strong inhibitory effect on classification.

The role of the V3A and the MT in the classification of the Random Motion state seems clear – both regions have been historically linked to motion perception and spatial awareness, crucial for tasks that require processing of dynamic, non-goal-directed visual stimuli (Pitzalis et al., 2010; Siman-Tov et al., 2022).

The superior temporal sulcus is a key region for higher-order social cognition and the integration of audiovisual social cues, helping to interpret socially relevant information (Erickson et al., 2017). Increased activation in the STS was associated with a decreased likelihood of classifying a trial as involving random motion — suggesting that when movement lacks social relevance, the demand for social perceptual processing in the STS is reduced.

Similarly, the IP0 region of the posterior parietal cortex, involved in multisensory integration, attention, and spatial processing, showed an inhibitory effect. This may indicate diminished engagement of integrative resources when the observed motion is meaningless.

Regarding the role of V3 as an inhibitor, it is important to note that while V3 and V3A are adjacent visual areas, V3A is more specialized for processing global motion and integrating visual information over larger fields, making it functionally distinct from V3 (Pitzalis et al., 2010). Thus, suppression of V3 may indicate that basic feature processing is less relevant during random motion, whereas more complex motion integration (as in V3A) is selectively engaged depending on task demands.

#### Motor Task: Right Hand or Foot

During this task participants were presented with visual cues that asked them to either tap their left or right fingers, or squeeze their left or right toes. Since all participants were right handed, the contralateral activation of motor regions was expected during the right hand or foot state and vice versa.

As expected, the contralateral motor regions 5L and 6mp carried the highest positive weight on classification. While the ipsilateral somatosensory area 1 exhibited a strong negative weight on classification.

Here, area 5L (part of the superior parietal lobule) is involved in integrating somatosensory and proprioceptive information for limb position and movement planning (Rech et al., 2019). However, area 6mp (part of the supplementary motor area), is critical for motor planning and execution (Hardwick et al., 2013). The involvement of both regions is expected due to the task’s directive motor nature and laterality.

The inhibitory role of the right area 1 on classification can be explained by consistently observed reduced involvement of the ipsilateral sensory cortex during dominant unilateral motor tasks, consistent with interhemispheric inhibition mechanisms that optimize contralateral processing.

#### Motor Task: Left Hand or Foot

In opposition to the Right Hand or Foot, the Left Hand or Foot condition involved a wide network of distributed contralateral and ipsilateral regions, acting both as activators and inhibitors of the classification.

The right (area 1, area 2) and left somatosensory regions (PFcm and PF), a left pre-motor region (6d), and a right opercular region (OP1) – all contributed positively to classification of the Left Hand or Foot state.

In contrast, the left primary motor cortex (area 4) carried the strongest negative weight, acting as the main inhibitor. In addition, adjacent contralateral motor areas - 5L, 5m and 6mp - were indicated as classification inhibitors. Interestingly, homologous to the activators somatosensory regions (right PFcm and PFt) were also found to decrease the likelihood of classifying the Left Hand or Foot state.

These results can be explained by the increased demands for controlling non-dominant limbs and compensatory recruitment of ipsilateral and contralateral networks. It has been repeatedly described that movements of the non-dominant left hand or foot in right-handed participants often elicit greater and/or bilateral activation in motor-related brain areas (Kim et al., 1993; Newton et al., 2005; Verstynen et al., 2005). In addition, the left motor task recruited a wider network of brain regions, which helped integrate somatosensory feedback to ensure accurate movement execution (Quirmbach & Limanowski, 2022). Here, the involvement of the complementary OP1 region, which is connected to but is not included in the motor or somatosensory cortices, underscores once again the greater task load for right-handed participants.

## 5 Conclusion

We demonstrated the relevance and applicability of classical ML algorithms for fMRI data classification, particularly in scenarios with limited data and a strong need for interpretability. All classical classification methods performed well, achieving high accuracy—especially for motor and language processing tasks. For some categories, the accuracy reached nearly 100%, highlighting the distinctiveness of the associated brain activity patterns.

A key focus of our study was identifying significant brain regions whose activity strongly correlated with specific brain states. We found unique sets of regions for each state, with minimal overlap between classes (confirmed by Jaccard coefficient analysis). This supports the hypothesis that each brain state is characterized by a distinct activity pattern arising from specific neural network dynamics. Notably, these regions align with established neuroscientific knowledge, validating our methodology’s ability to identify neural correlates of cognitive states.

Further analysis revealed that high-accuracy states required relatively few significant regions, suggesting focal neural signatures. In contrast, lower-accuracy states involved more distributed activations. In such cases, it was not possible to identify a distinct feature set sufficient for accurate classification, which may be due to these states being formed by the joint activity of multiple brain regions. This result highlights the importance of further investigation into such states to identify their characteristic activation patterns.

The results obtained can be used for further exploration of the neurobiological mechanisms underlying cognitive processes. The proposed approach represents an effective and accessible tool for a broad group of scientists to analyse both spatial and temporal dynamics of brain activity, identifying significant regions and their interactions. It is a valuable method for studying the neural basis of cognitive processes and the functional organization of the brain, especially in scenarios with limited data.

In this second part of our study, our primary goal was to demonstrate, using rigorous mathematical methods, the importance of temporal dynamics in the functional characterization of significant brain regions identified by linear classifiers. The results across all three analyses consistently support this conclusion.

First, the correlation analysis showed that top-ranked brain regions exhibit markedly stronger temporal correlations compared to low-ranked regions, and that these correlation distributions are separable with good accuracy. This indicates that discriminative brain states emerge from coordinated activity patterns across specific neural circuits.

Second, the temporal structure test confirmed that, for nearly all brain region pairs, the distributions of correlations obtained from original time series are statistically distinct from those derived from shuffled time series. This provides direct evidence that preserving temporal structure is critical for capturing meaningful functional interactions.

Taken together, these results demonstrate that the discriminative power of significant brain regions is closely tied to their temporal dynamics. Our analyses highlight that correlation-based measures consistently capture the functional importance of these regions, thereby validating their role in shaping distinct cognitive brain states.

In conclusion, the proposed approach has the potential to be applied in different neuroimaging settings and scenarios. For example, integration of our ML classification pipeline with naturalistic stimuli paradigms, such as movie watching or storytelling tasks, could enhance ecological validity and classification accuracy, as naturalistic fMRI has shown superior predictive power compared to traditional task-based approaches (Zhang et al., 2021). Furthermore, our approach could serve as the principled feature selection for multimodal integration with structural MRI and diffusion tensor imaging, providing interpretable solutions, for instance, in clinical cases (Odusami et al., 2024). Finally, combining our pipeline with advanced nonlinear models could further refine the mapping between brain activity patterns and cognitive states, while preserving the interpretability achieved through the present linear analysis.

## Code availability

The source code and data are available at: https://github.com/valeria-kirova/HCP-analysys.

